# Enhancing CYP450-Ligand Binding Predictions: A Comparative Analysis of Ligand-Based and Hybrid Machine Learning Models

**DOI:** 10.1101/2025.06.05.658010

**Authors:** Anastasiia Tikhonova, Eric Chun Yong Chan, Hao Fan

## Abstract

Predicting cytochrome P450 (CYP450) ligand binding is critical in early-stage drug discovery, as CYP450-mediated metabolism profoundly influences drug efficacy, safety, and adverse reaction risks. However, experimental determination of CYP450-ligand interactions remains resource- and time-intensive, underscoring the need for robust computational alternatives. While ligand-based methods are commonly employed, they often fail to fully account for structural intricacies governing protein-ligand interactions. To address this gap, we developed a hybrid machine learning framework integrating ligand descriptors, protein descriptors, and protein-ligand interaction descriptors, that include molecular docking-derived parameters, rescoring function components from multiple algorithms and structural interaction fingerprints (SIFt). Evaluated on CYP1A2 and CYP17A1 isoforms, our model demonstrated superior predictive accuracy in cross-validation compared to stand-alone molecular docking and ligand-based approaches. Furthermore, benchmarking against state-of-the-art tools —SwissADME and ADMETlab 3.0 — revealed enhanced performance in binding prediction. This work establishes a versatile framework for advancing computational tools to prioritize CYP450 binding assessments during drug discovery.

## INTRODUCTION

Cytochrome P450 (CYP450) enzymes catalyze the metabolism of xenobiotics and endogenous compounds, playing a key role in homeostasis and drug detoxification. Five key isoforms — CYP1A2, CYP2C9, CYP2C19, CYP2D6, and CYP3A4 — mediate approximately 75% of drug metabolism in humans [1]. Their broad substrate specificity and conformational flexibility, particularly within the heme-associated binding region, enable metabolism of chemically diverse compounds [2,3]. However, this versatility complicates binding prediction, as CYP450 enzymes interact with structurally heterogeneous ligands, posing challenges for drug design [4]. Accurate CYP450-ligand binding prediction methods are therefore crucial for optimizing drug candidates.

Experimental techniques such as high-throughput screening (HTS) and X-ray crystallography have deepened the understanding of CYP450-ligand binding and served as a foundation for computational methods. However, these methods are resource-intensive, requiring specialized instrumentation, substantial time, and technical expertise [5]. Physics-based computational methods, including molecular docking and molecular dynamics (MD) simulations, are widely used to study CYP450-ligand interactions. Docking methods can predict ligand orientation and estimate binding energy, while MD simulations can provide detailed insights into the dynamic behavior of protein-ligand complexes. However, these methods exhibit certain limitations. Docking relies on high-resolution crystal structures, which are unavailable for some CYP450 isoforms. Moreover, docking methods face inherent challenges in accurately modeling induced-fit effects and the structural flexibility of CYP450 binding sites, which can accommodate diverse ligands, including large macrolides [6,7]. While MD simulations address these limitations by accounting for conformational dynamics, their high computational cost often precludes their application in large-scale virtual screening efforts.

To address the limitations of physics-based computational methods, ligand-based computational methods provide an alternative for binding prediction without requiring structural data of the protein. On the other side, ligand-based computational methods frequently encounter significant dataset imbalances due to the systematic preference for publishing data on CYP450-binding molecules over non-binding molecules. Such imbalances can compromise model performance by leading to overfitting due to limited representation of non-binding molecules, potentially reducing predictive accuracy for novel ligands with distinct activity profiles [8]. These ligand-based approaches, while valuable, do not explicitly capture protein-specific characteristics, such as conformational flexibility or interactions with multiple binding regions, which may limit their applicability in complex scenarios.

To overcome these limitations, hybrid machine learning (ML) models have been developed to integrate ligand descriptors and protein-ligand interaction fingerprints. For example, Katsila et al. introduced such a model that incorporates RDKit ligand descriptors alongside Protein-Ligand Extended Connectivity (PLEC) fingerprints to account for a more nuanced representation of CYP450-ligand binding [9]. Based on their findings, the authors concluded that a robust framework with multiple feature types is needed for CYP450-ligand binding prediction to accommodate structural variability in CYP450 isoforms [9].

In this study, we systematically evaluated CYP450-ligand binding prediction models by investigating the following questions: 1) How does integrating protein-specific descriptors and protein-ligand interaction features with ligand-based descriptors influence ligand prediction accuracy? 2) Does a weighted ensemble machine learning framework outperform individual models in predicting CYP450-ligand binding? 3) Does incorporating multiple CYP450 protein structures enhance model performance compared to reliance on a single protein structure?

By addressing these questions, we developed a hybrid framework that integrates ligand-based descriptors, protein structural features, and protein-ligand interaction fingerprints. This multi-faceted approach allows us to capture both chemically diverse properties of ligands and the structural dynamics of CYP450 isoforms, thereby enabling a holistic characterization of binding mechanisms. This framework can be used to streamline drug discovery processes and improve safety assessments associated with CYP450-mediated metabolism. Furthermore, it establishes a foundation for generalizable, structure-informed predictive models that address limitations in current in-silico approaches for CYP450-ligand binding.

## RESULTS AND DISCUSSION

### Molecular Docking Performance as a Baseline

Molecular docking, as a stand-alone approach, demonstrated limited predictive capability across four CYP450 isoforms (CYP2C9, CYP2C19, CYP2D6, and CYP3A4), as shown in Supplementary Table 1. In this work, molecular docking performance was assessed using ligand enrichment (logAUC), which can be described as the area under the curve of enrichment plot, with x axis in logarithmic scale [10]. For CYP2C9, the three crystal structures (PDB code: 5A5I, 1OG5, 6VLT) yielded logAUC values of 0.24, 0.26, and 0.24, respectively. For CYP2C19, the crystal structure (PDB code: 4GQS) and the RoseTTAFold model achieved logAUC of 0.24 and 0.23, respectively, while the AlphaFold model showed improved performance (logAUC=0.36). For CYP2D6, all three structures (PDB code: 3TBG, 4XRZ, and 3QM4) showed comparable performance with logAUC of 0.24, 0.24, and 0.23, respectively. Similarly, CYP3A4 structures (PDB code: 4D6Z, 2J0D, and 8EXB) also showed comparable performance with logAUC of 0.24, 0.25, and 0.24, respectively.

To further evaluate the generalizability of docking, we applied the same approach to two isoforms that were set aside for machine learning validation. For CYP1A2, molecular docking using the crystal structure (PDB code: 2HI4) yielded logAUC value of 0.20, while docking against the AlphaFold and RoseTTAFold models achieved logAUC values of 0.26 and 0.22, respectively. CYP17A1 structures (PDB code: 6WR1, 3SWZ, and 8FDA) demonstrated comparable performance with logAUC values of 0.24, 0.24, and 0.23, respectively. While showing moderate performance, these results highlight the limitation of structure-based molecular docking method in handling interactions between CYP450 proteins and diverse ligands.

In this study. consensus enrichment was also calculated for each protein, by combining docking results of the 3 structures/models. For each docked compound (ligands and decoys), the best docking score across all structures/models was used for the ranking of that compound [11]. The consensus enrichment value is 0.30, 0.35, 0.26, 0.25, 0.28, 0.25 for CYP2C9, CYP2C19, CYP2D6, CYP3A4, CYP1A2, and CYP17A1, respectively, better than or comparable to the best performing single structure/model in all cases. These findings suggest that incorporating multiple structures/models in docking screens can offers better prediction for CYP450 ligands than using a single structure, especially in real-world applications where it is difficult to know the best-performing structure for novel ligands a prior. In the following development of machine learning models, we also explored the effect of including multiple protein structures.

### Protein and Protein-Ligand Descriptors Improve Performance of Hybrid Models

To evaluate the impact of incorporating information on protein and protein-ligand interactions, we compared the performance of ligand-based models (using only ligand descriptors) against hybrid models (combining ligand descriptors with protein descriptors and protein-ligand interaction descriptors, that include molecular docking-derived parameters, rescoring function components and SIFt) across both the validation and external testing isoforms (the workflow design is presented in Figure 1).. For simplicity, we first focused on the top scored machine learning model (without weighted ensemble framework) to isolate the contribution of individual feature sets.

**Figure 1.**
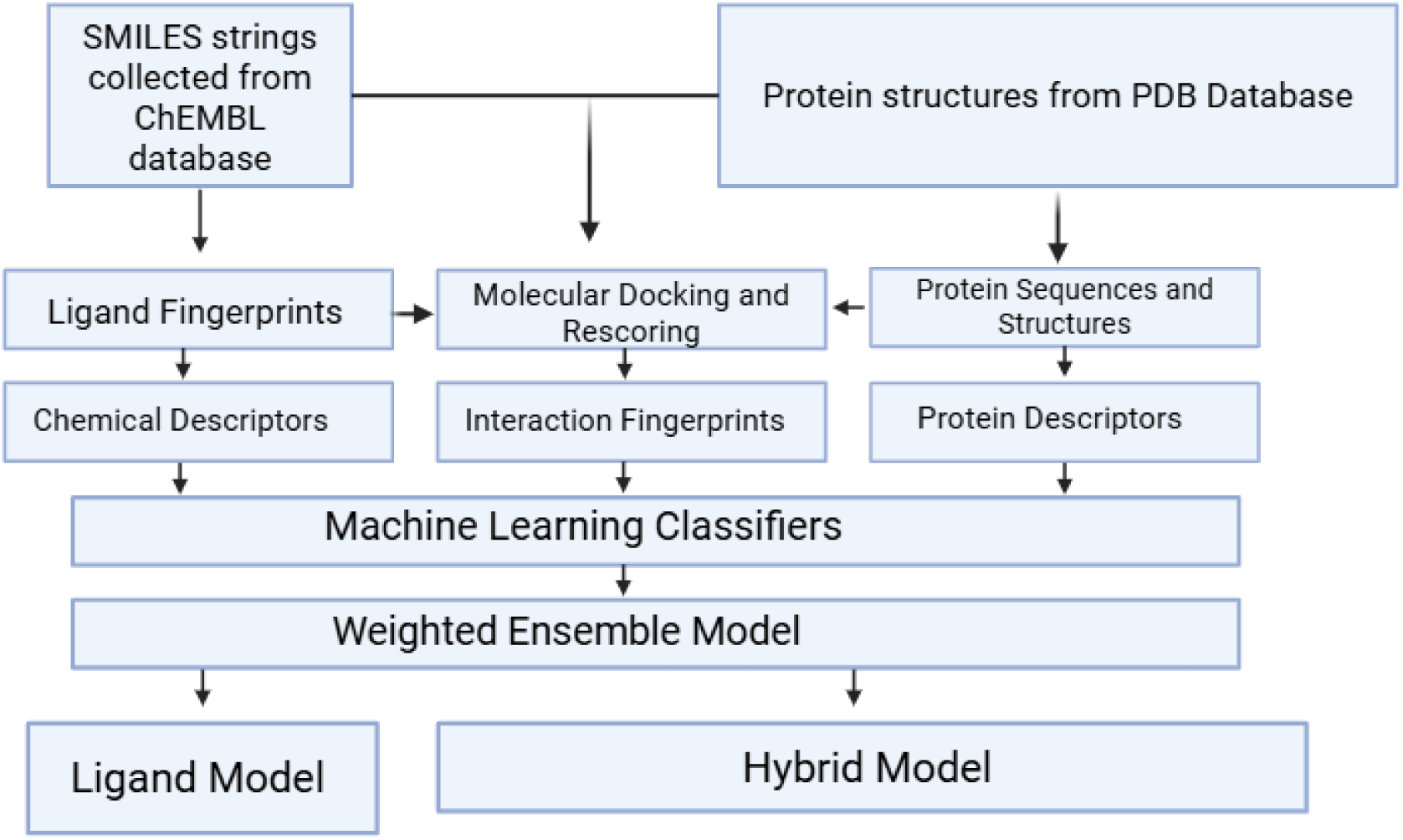
Outline of the Workflow for the Proposed Machine Learning Framework. Created in BioRender. Tikhonova, A. (2025). https://BioRender.com/rwcbj4k

The performance of the top-scored ligand-based model (L_TS) across isoforms was consistent but limited, achieving logAUC values of 0.40 ± 0.022 for CYP2C9, 0.38 ± 0.016 for CYP2C19, 0.45 ± 0.043 for CYP2D6, and 0.38 ± 0.016 for CYP3A4 in cross-validation (Figure 2). The top-scored hybrid model including one single protein structure (H_TS_S) demonstrated significantly improved performance in cross-validation with logAUC values of 0.90±0.020 for CYP2C9, 0.81 ± 0.039 for CYP2C19, 0.90±0.059 for CYP2D6, and 0.97±0.074 for CYP3A4, highlighting the synergistic effect of incorporating protein descriptors and protein-ligand interaction descriptors (Supplementary Table 2). This correlates with the findings from the feature importance analysis, which will be discussed in subsequent sections of this article, where protein-ligand interaction features such as the Glide Docking Score, ITScoreAff scoring function, and SIFt were identified as dominant predictors. These descriptors could contribute to the improved performance of the CYP450-ligand binding prediction, as they capture essential interactions — such as electrostatic and hydrophobic interactions — that directly influence ligand binding to CYP450s. These results highlight the importance of combining ligand-based descriptors with protein and protein-ligand interaction descriptors to capture the full complexity of CYP450 ligand binding. The variability in standard deviation across isoforms can be attributed to the stochastic nature of automated machine learning approaches and the inherent diversity of molecules within the dataset.

**Figure 2.**
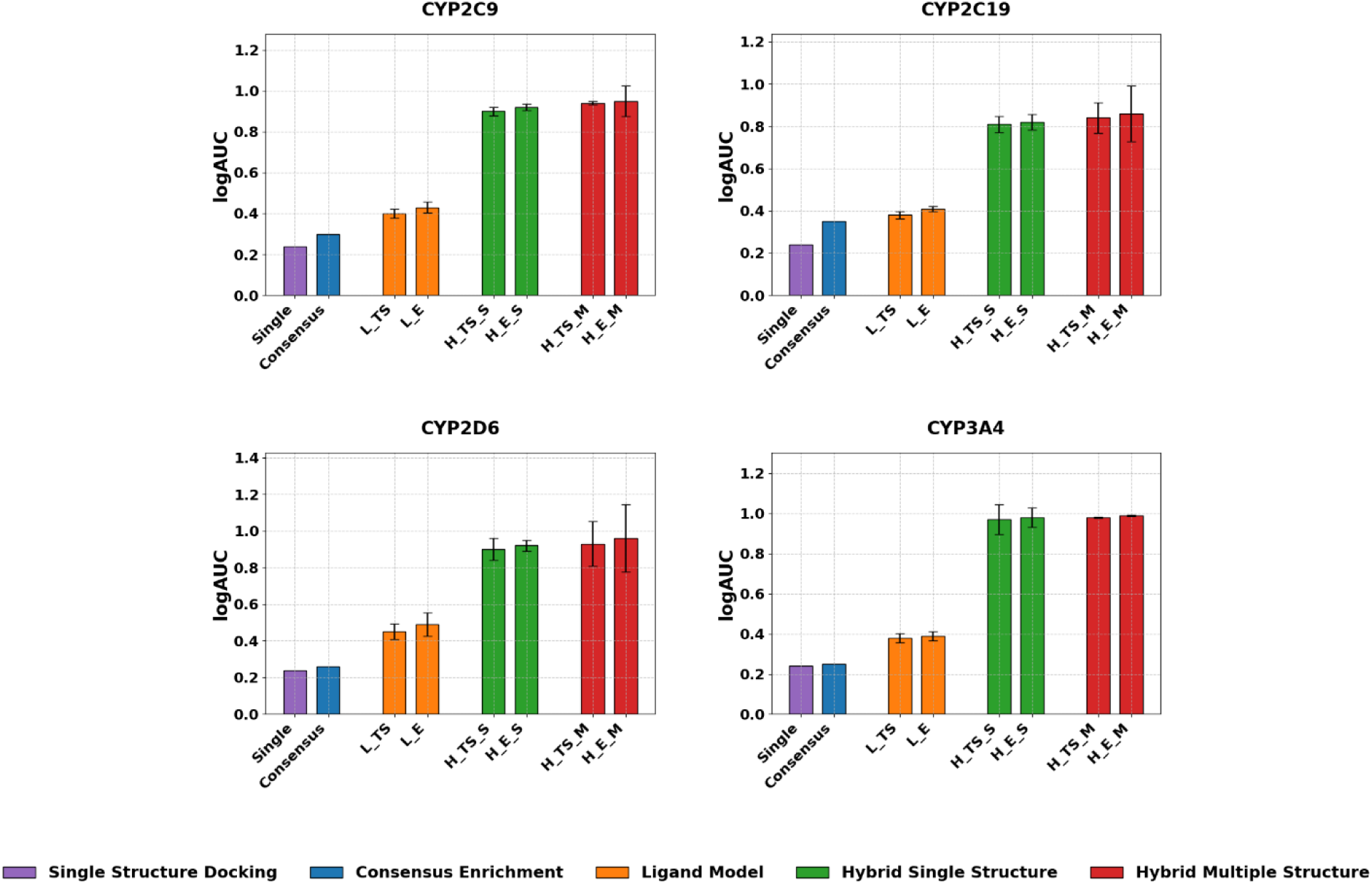
Predictive performance comparison of docking, ligand-based, and hybrid models across CYP2C9, CYP2C19, CYP2D6 and CYP3A4 isoforms (represented by logAUC values). Error bars indicate standard deviation across 50-fold cross-validations. Single Structure Docking Enrichment (in purple): logAUC values are based on molecular docking scores from the highest resolution crystal structure of the protein. Consensus Enrichment (in blue): logAUC values are based on the most negative docking score across three selected structures of the protein. Ligand Model (in orange): top scored model trained using ligand descriptors (L_TS) and weighted ensemble model trained using ligand descriptors (L_E). Hybrid models trained on ligand, protein and protein-ligand interaction descriptors (including molecular docking-derived parameters, rescoring function components and SIFt) of single protein structure (green): top scored model (H_TS_S) and weighted ensemble model (H_E_S); and on multiple protein structures (in red): top scored model (H_TS_M) and weighted ensemble model (H_E_M).

### Weighted Ensemble Model Demonstrates Marginal but Consistent Performance Gain Over Single Top Scored Model

Among ligand-based models, the performance of the top scored model is slightly and consistently improved across all isoforms when weighted ensemble approach was applied (Figure 2). For CYP2C9, the weighted ensemble ligand model (L_E) achieved a logAUC of 0.43 ± 0.027 compared to the logAUC of 0.40 ± 0.022 from the top scored model (L_TS). Similar moderate improvements were observed for CYP2C19 (0.41 ± 0.012 vs. 0.38 ± 0.016), CYP2D6 (0.49 ± 0.062 vs. 0.45 ± 0.043), and CYP3A4 (0.39 ± 0.022 vs. 0.38 ± 0.023), respectively (Supplementary Table 2). These limited improvements suggest top-performing models already capture most relevant ligand-based interactions, with weighted ensemble approach providing marginal benefits by averaging noise and incorporating diverse feature representations.

Similarly, the weighted ensemble approach also brought improvement to hybrid models (Figure 2). The weighted ensemble hybrid model (H_E_S) consistently outperformed the top scored hybrid model (H_TS_S) across all isoforms: CYP2C9 (0.92 ± 0.015 vs. 0.90 ± 0.020), CYP2C19 (0.82 ± 0.038 vs. 0.81 ± 0.039), CYP2D6 (0.92 ± 0.029 vs. 0.90 ± 0.059), and CYP3A4 (0.98 ± 0.048 vs. 0.97 ± 0.074). Again, while these 1-2% improvements demonstrate some benefit of weighted ensemble approach in modelling CYP450-ligand interactions, these small differences suggest that the top-scored model may already capture the majority of relevant interactions.

### Incorporating Multiple Protein Structures Enhances CYP450 Ligand Binding Prediction Accuracy

The weighted ensemble hybrid model including multiple protein structures for each isoform (H_E_M) yielded additional improvements in ligand enrichment on top of the weighted ensemble hybrid model including only one single protein structure for each isoform. The enhancement of logAUC values ranges from 1 to 4%, for CYP2C9 (0.95 ± 0.074 vs. 0.92 ± 0.015), CYP2C19 (0.86±0.131 vs. 0.82±0.038), CYP2D6 (0.96±0.183 vs. 0.92±0.029) and CYP3A4 (0.99 ± 0.004 vs. 0.98 ± 0.048), respectively.

The larger standard deviation observed in some isoforms, particularly CYP2D6 and CYP2C19, can be attributed to the variability introduced by AutoGluon’s dynamic parameter tuning, which adjusts hyperparameters based on isoform-specific data characteristics. This reflects AutoGluon flexibility but also highlights the challenges of achieving consistent optimization across structurally diverse CYP450 isoforms. The diversity of structural structures included in the multi-structure models enhances robustness but might potentially introduce parameter-specific noise. Additionally, aside from Autogluon stochastic nature, there might be other reasons for particularly high standard deviations in CYP2C9 and CYP2D6 isoforms, including the limited size of CYP2C19 ligand dataset (149 ligands after clustering —the smallest among all isoforms), which reduces model robustness and amplifies the effects of sampling variability during cross-validation. Additionally, only one experimental crystal structure was available for CYP2C19; the remaining two protein structures were generated by AlphaFold and RoseTTAFold predictions, potentially introducing structural noise into docking-derived descriptors and contributing further to prediction instability. For CYP2D6, the variability might be caused by the choice of crystal isoforms – while the two structures 4XRZ and 3QM4 have an active site that forms 16 contacts on average between the crystal ligand and the protein (estimated by SIFt “Any Contact” descriptor obtained from Glide docking poses), the other structure 3TBG was determined in a more closed conformation, resulting in fewer contacts (with average being 3). Considering this, future method development may consider alternative optimization strategies to mitigate the impact of diverse protein structures, ensuring that specific protein isoforms do not disproportionately influence the model’s learning.

### Evaluating model generalizability by external testing on isoforms CYP1A2 and CYP17A1

To evaluate the ability of the model to generalize beyond the training data, we performed external testing on CYP1A2 and CYP17A1 isoforms that were not included in the training set, assessing the predictive performance on unseen protein targets. Similar to the cross–validation described in the previous sections, we assessed the impact of incorporating protein and protein–ligand interaction features, the weighted ensemble method, and multiple protein structures on the model’s predictive performance.

First, to evaluate the impact of incorporating protein and protein–ligand interaction descriptors to the model in external testing, we compared the performance of the top scored ligand model (L_TS) to the top scored hybrid model (H_TS_S) on the CYP1A2 and CYP17A1 isoforms. The hybrid model (H_TS_S) improved prediction accuracy for both CYP1A2 (logAUC of 0.31 vs. 0.28 for ligand-based model, representing a 1.1-fold improvement) and CYP17A1 (logAUC of 0.95 vs. 0.47 for ligand-based model, a 2-fold increase in prediction accuracy). While the improvement for CYP1A2 is marginal, the remarkably improved logAUC for CYP17A1 suggests that the hybrid model may offer greater predictive power for certain isoforms (Supplementary Table 3). This difference in performance could be explained by the fact that CYP17A1 ligands are more chemically similar to those in the training set, with over 90% of them having a maximal Tanimoto similarity of 0.70 or above to the training set, while only 54% of CYP1A2 ligands exhibit such similarity, making CYP17A1 a more favorable target for both the ligand-based and hybrid models. Considering this fact, the ligand similarity between training set isoforms and testing set isoforms should be considered for building similar frameworks. To investigate other potential differences in logAUC for CYP17A1 and CYP1A2, we calculated the mean values of ligand, protein and protein-ligand interaction descriptors for each isoform and compared the obtained mean values between CYP17A1 and CYP1A2 (CYP17A1-CYP1A2). The most significant differences in mean values were observed in descriptors derived from molecular docking: Glide Emodel, an empirical scoring function that has a more significant weighting of the force field components compared to Glide Score, showed the largest difference (Δ = 42.29), followed by ITScoreAff rescoring function (Δ = 33.21). These differences suggest that ligand binding to CYP17A1 is associated with a lower energy score compared to CYP1A2. The additional structural descriptors derived from docking are likely to provide a more accurate representation of the binding characteristics for this isoform. Next, to assess the potential impact of using weighted ensemble approach, we compared the top scored model to weighted ensemble model on CYP1A2 and CYP17A1 isoforms. Weighted ensemble models also showed a marginal improvement in external testing, with weighted ensemble hybrid model (H_E_S) only one single protein structure achieving higher logAUC values for CYP1A2 (0.36 vs. 0.31) and CYP17A1 (0.97 vs. 0.95) compared to top scored model (H_TS_S).

To evaluate the impact of incorporating multiple protein structures, we performed external testing using multiple structures (Figure 3). Notably CYP1A2 has just one crystal structure, with the other two structures being AlphaFold and RoseTTAFold models. Hybrid weighted ensemble model incorporating multiple structures (H_E_M) improved performance in external testing for CYP1A2 (0.47 vs. 0.36) and CYP17A1 (0.98 vs. 0.97) compared to their single structure counterparts (H_E_S). These results suggest that the usage of multiple structures for CYP450-ligand binding prediction frameworks may offer benefits primarily for certain challenging isoforms with specific structural characteristics.

**Figure 3.**
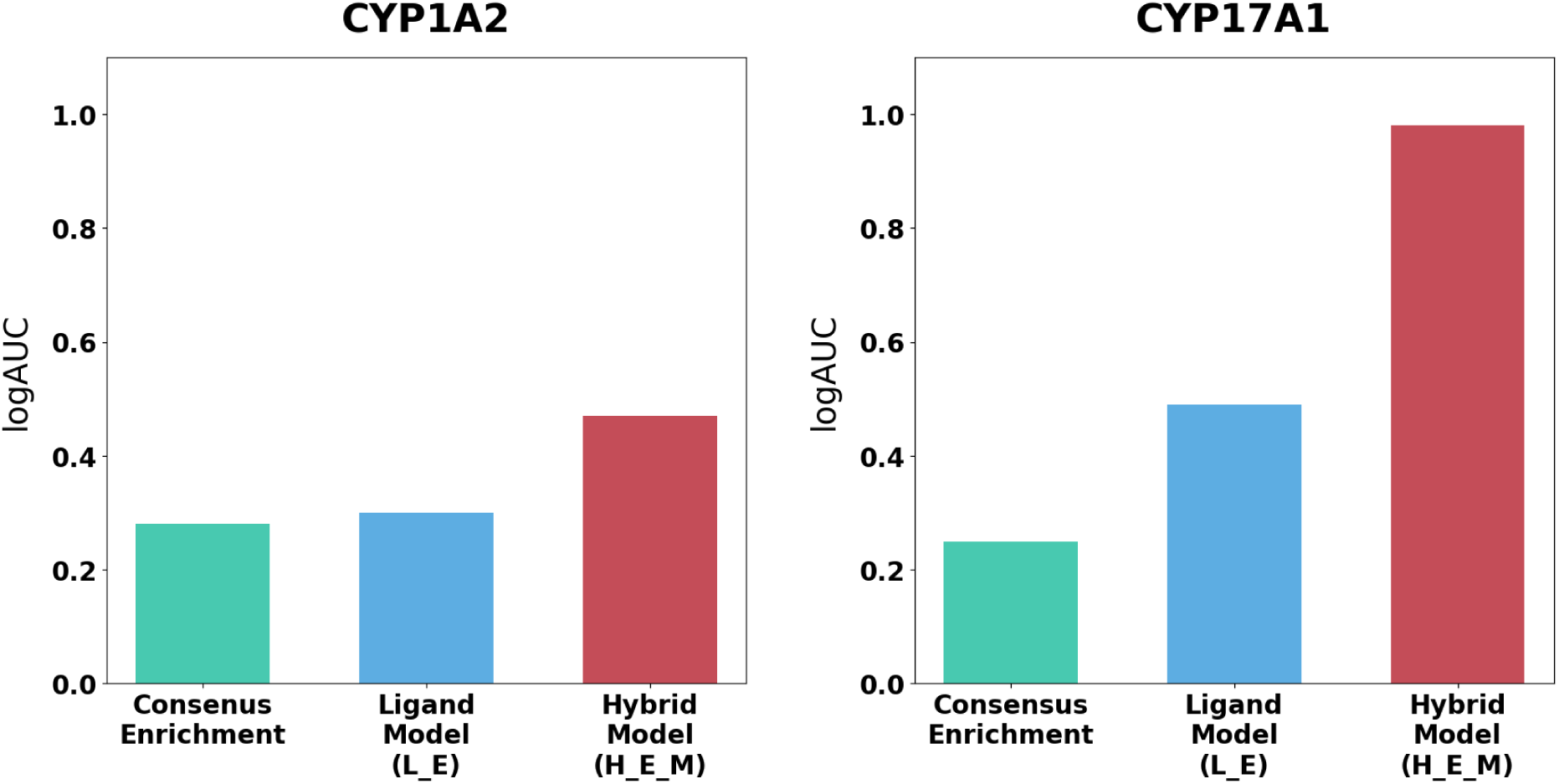
External testing results comparing consensus enrichment, weighted ensemble ligand model (L_E), and weighted ensemble multiple structure hybrid model (H_E_M) performance in CYP1A2 and CYP17A1 isoforms.

### Evaluating Feature Importance in CYP450-Ligand Binding Predictions

The feature importance analysis was conducted for the weighted ensemble multiple structure hybrid model (H_E_M) which demonstrated the best prediction accuracy, so it can serve as the representative for understanding the main factors (top 10) impacting prediction performance, ensuring that the analysis highlights the key contributors for accurate predictions (Figure 4). The list of all used ligand, protein and protein and protein-ligand interaction features is given in Supplementary Table 4.

**Figure 4.**
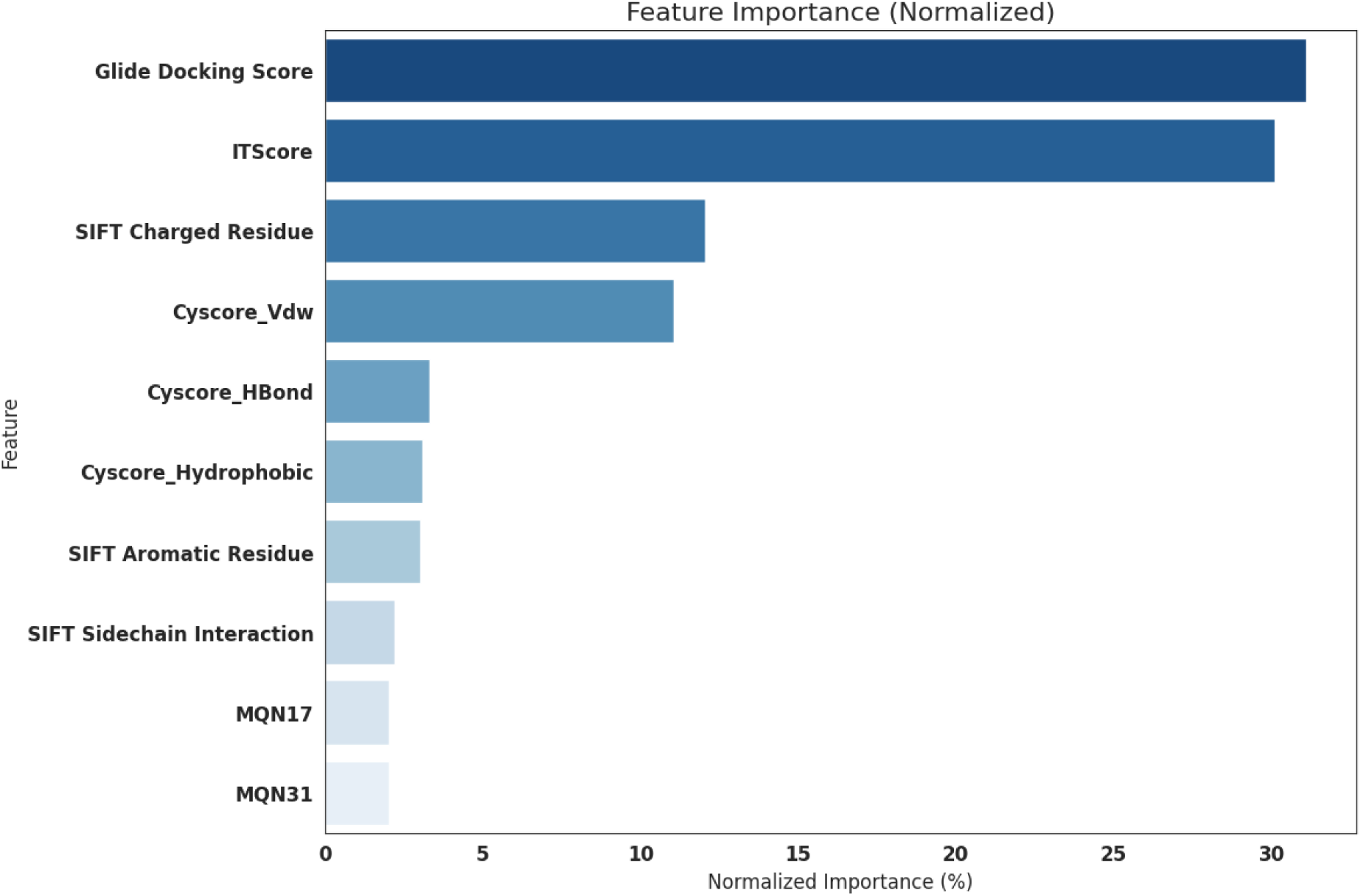
Relative importance of molecular descriptors and interaction features in the Weighted Ensemble Multiple Structure Hybrid Model for CYP450 binding prediction. Features are shown as normalized importance percentages.

Descriptors associated with molecular docking emerged as the dominant predictors, with Glide Docking Score (0.31) and ITScoreAff scoring function (0.30) contributing the most to the model’s performance (Supplementary Table 5). Collectively, these two features constitute roughly 61% of the total feature importance, underscoring the major role of descriptors that capture CYP450-ligand binding properties.

Protein-ligand interaction fingerprints formed the second tier of influential features. SIFt Charged Residue interactions descriptor that represents the number of all charged interactions in the CYP450-ligand complex, showed moderate importance (0.12), suggesting that ionic and dipolar interactions substantially modulate binding thermodynamics. Several descriptors in this category were derived from Cyscore, an empirical scoring function that predicts protein–ligand binding affinity by combining hydrophobic, van der Waals, and hydrogen-bond energies with ligand entropy, using a curvature-weighted surface area model to more accurately capture hydrophobic effects [12]. Cyscore_Vdw, a descriptor derived from the Cyscore scoring function that captures the Van der Waals interaction energy within the complex, also contributed notably (0.11) — underscoring the importance of van der Waals forces and steric complementarity in CYP450– ligand binding. The remaining features, while contributing to a lesser extent, still captured important aspects of protein–ligand interactions: Cyscore_HBond (0.033), reflecting hydrogen-bonding interactions in docking poses; Cyscore_Hydrophobic (0.031), representing the hydrophobic interactions of the complex; and SIFt Aromatic Residue (0.030), indicating the presence of aromatic stacking. Altogether, these descriptors highlighted distinct facets of the binding landscape. These descriptors indicate essential stabilizing interactions within the CYP450 active sites, including hydrogen bonding, hydrophobic contacts, and π-stacking with aromatic residues. SIFT Sidechain Interaction (0.022), reflecting the number of sidechain contacts in the complex, and molecular fingerprint descriptors (MQN17 and MQN31, representing cyclic double bonds and trivalent nodes count, both 0.02), showed lower importance, suggesting that while ligand structural properties and sidechain contacts contribute to binding prediction, they play a more supplementary role compared to the energy functions and key interaction patterns.

This hierarchical distribution of feature importance highlights that accurate CYP450-ligand binding prediction relies primarily on comprehensive description of protein-ligand physical interactions, with specific interaction types providing additional refinement. The significant contribution of Glide Docking Score and ITScoreAff to predictive performance emphasizes the value of incorporating binding energy scores to CYP450-ligand binding prediction models.

### Benchmarking with Existing Models for CYP450-Ligand Binding Prediction

The benchmarking analysis was conducted to evaluate the performance of the Weighted Ensemble Multiple Structure Hybrid Model (H_E_M) against two state-of-the-art CYP450-ligand binding prediction tools, SwissADME and ADMETlab 3.0 [13,14], both of which rely on ligand descriptors without explicitly accounting for either protein descriptors or protein-ligand interaction descriptors.

Our selection of CYP3A4 and CYP1A2 isoforms for this benchmarking analysis was based on several factors: 1) the two established models SwissADME and ADMETlab contain these two isoforms; 2) these two isoforms cover different conditions regarding protein structure availability — CYP1A2 has only one crystal structure available, while CYP3A4 has multiple crystal structures, allowing us to assess the impact of relying solely on crystal structures versus incorporating structures from AlphaFold and RoseTTAFold; 3) CYP3A4 was included in our training set, whereas CYP1A2 was held out for external testing, enabling us to evaluate both the model’s fitting capabilities and its generalizability to novel isoforms.

The comparison focused on two key evaluation metrics: AUPR (Area Under the Precision-Recall Curve) and AUC (Area Under the ROC Curve) scores, which are reported in the majority of machine learning works focusing on CYP450-ligand binding prediction and provide standard reference points for benchmarking. Benchmarking was performed using binders and non-binders from the ChEMBL database that were reported after the publication of SwissADME and ADMETlab reference methods. These ligands were not included in our training dataset nor in the training data of the other models used in benchmarking, ensuring a fair evaluation of each method’s predictive ability on novel chemical entities. To ensure structural diversity, the obtained ligands were clustered using the Butina method with a Tanimoto similarity threshold of 0.6, and the cluster centroids were chosen for benchmarking.

The Hybrid Model demonstrated better performance for both CYP isoforms, (CYP3A4: AUC = 0.76, AUPR = 0.78; CYP1A2: AUC = 0.61, AUPR = 0.57) than both ADMET 3.0 (CYP3A4: AUC = 0.47, AUPR = 0.51; CYP1A2: AUC = 0.44, AUPR = 0.47) and SwissADME (CYP3A4: AUC = 0.51, AUPR = 0.53; CYP1A2: AUC = 0.56, AUPR = 0.54) (Figure 5, Supplementary Table 6). The Hybrid Model’s improved predictive performance can be attributed to its ability to integrate ligand, protein and protein-ligand interaction descriptors (including molecular docking, rescoring and SIFt), which allows for a more comprehensive representation of the CYP450-ligand binding. While CYP3A4 was included in our training set and CYP1A2 was held out for external testing, the mean maximum pairwise similarity between the benchmarking compounds and the training dataset was comparable for both CYP3A4 (0.32 ± 0.096) and CYP1A2 (0.28 ± 0.048). The new ligand set was not included in the initial training or external testing and was collected from the new entries to ChEMBL database, which explains the structural difference from the testing set. The factor contributing to the difference in model performance between CYP1A2 and CYP3A4 may be the structural characteristics of the binding pocket. Literature reports have suggested that CYP1A2 features a narrower and more constrained binding site, which can limit the diversity of ligand interactions and reduce the effectiveness of protein–ligand interaction features [15, 16, 17]. To validate these observations, we performed binding pocket analysis using the PockDrug statistical model [18]. Our analysis confirmed that CYP1A2 has a smaller average binding pocket volume (Volume Hull = 5748 ± 791 Å³) compared to CYP3A4 (Volume Hull = 8082 ± 3972 Å³). The larger and more flexible binding environment of CYP3A4 likely facilitates a wider range of ligand accommodations and interactions, enabling the hybrid model to better leverage docking-derived features and resulting in a more substantial improvement in predictive performance.

**Figure 5.**
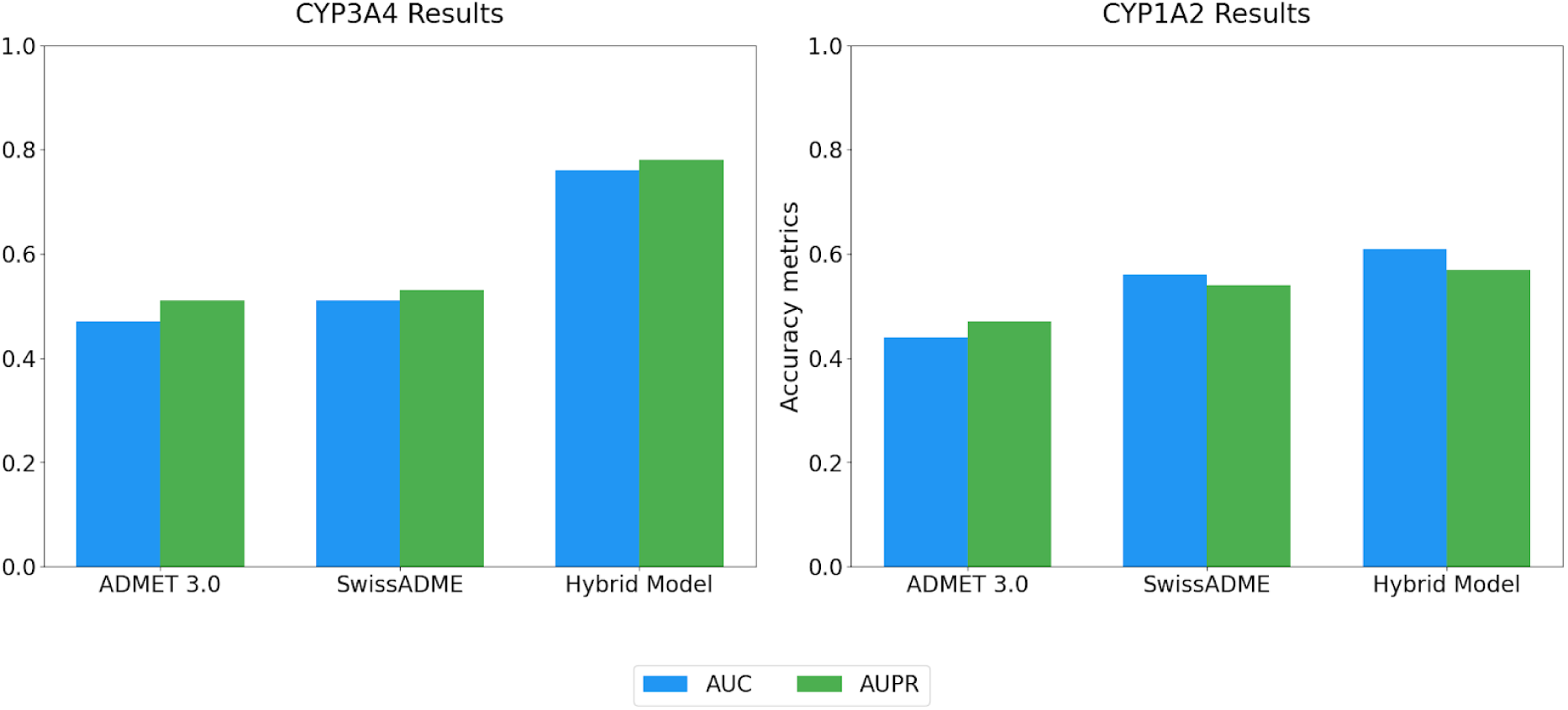
Model Performance Comparison between Hybrid Model, ADMET 3.0, and SwissADME across two key metrics: AUC and AUPR for CYP3A4 and CYP1A2 isoforms.

## CONCLUSION

Despite the availability of existing prediction methods for CYP450-ligand binding, significant opportunities exist for improving predictive accuracy through integration of recent advances in machine learning architectures and implementing more specific descriptors. In this study, we aimed to quantitatively measure the effects of integrating protein descriptors and protein-ligand interaction descriptors (including molecular docking, rescoring and SIFt) with ligand-based descriptors, during CYP450-ligand binding predictions. Our systematic evaluation demonstrated that incorporating protein and protein-ligand interaction descriptors, utilizing weighted ensemble approach and including multiple structures of proteins contributed measurably to prediction performance compared to ligand-based models.

While ligand-based models achieved only moderate performance (logAUC 0.38-0.49), our hybrid approach significantly improved prediction accuracy (logAUC 0.86-0.99) across multiple CYP450 isoforms in cross-validation. This improvement highlights the importance of capturing the complete binding environment rather than focusing solely on ligand properties. Furthermore, our cross-validation results revealed that implementing a weighted ensemble approach consistently outperformed individual top-scoring models by ∼2% across all isoforms. This finding suggests that different model architectures capture complementary aspects of binding mechanisms, and their combination provides more robust predictions than any single model, regardless of its individual performance metrics. In addition, we observed that utilizing multiple protein structures for each CYP450 isoform, rather than a single representative conformation, enhanced prediction performance—up to ∼5% improvement in cross-validation. This improvement indicates that structural diversity plays a crucial role in accurately modeling the conformational flexibility inherent in CYP450-ligand interactions, capturing essential binding mechanisms that single-structure models frequently miss.

Two ligand-based machine learning models (SwissADME, ADMET3.0) for CYP450 binding prediction were compared to the obtained hybrid model (H_E_M) with the highest predictive accuracy in this work. Both tools have their place in the early stages of the drug discovery pipeline, particularly when multiple prediction tools need to be compared for consensus. The hybrid model showed improved performance metrics for the CYP17A1 isoform, indicating that incorporating structural information about protein-ligand interactions may provide additional predictive value beyond what can be achieved with ligand properties alone, particularly for specific CYP450 isoforms where binding mechanisms are more complex or dependent on protein conformational states.

We acknowledge that this study relies heavily on the ChEMBL database for collecting ligands that are used for building datasets, which may affect the generalizability of this model for novel described ligands that are highly dissimilar from the ligands that were used for training the models. In this study, we used DUD-E decoys to balance the dataset by compensating for the lack of negative binding results. However, while this is a common practice, it may not fully reflect the characteristics of real-world non-binders, potentially affecting the generalizability of the model. Moreover, the selection of only three conformations per isoform may not capture the full range of protein flexibility. We recommend that future workflow development could benefit from integrating more crystal structures of the proteins to incorporate more protein flexibility and including recently described ligands to improve model performance and generalizability in real-world scenarios. Furthermore, developing models accommodating genetic variants in CYP450 could introduce another layer of complexity and depth in predicting CYP450-ligand binding

## METHODS

### Dataset collection and Processing: Protein Selection

The isoforms CYP2C9, CYP3A4, CYP2C19, and CYP2D6 were selected as primary targets for model training, while CYP1A2 and CYP17A1 were held out for external testing. The experimentally resolved 3D structures of these proteins were retrieved from the RCSB Protein Data Bank (PDB) (Table 1) [19]. The protein structures selected for docking were required to meet several criteria. Specifically, they needed to be co-crystallized with a bound ligand, possess no mutations within their binding sites, and, where possible, have high-resolution X-ray crystallography data. In cases where mutations were present, the structures were reverted to their native forms using the Schrödinger Maestro Suite, following established correction protocols to ensure accuracy in binding site representation [15]. To comprehensively capture the structural diversity of CYP450-ligand binding pockets, three distinct protein structures were selected from the available PDB entries for each isoform, where possible. The first structure was chosen based on the highest resolution structure available in the PDB. The second structure was selected for having the biggest root-mean-square deviation (RMSD) of the binding site with respect to the first structure, ensuring the inclusion of significant backbone variations. The third structure was identified by maximizing the Tanimoto distance between the ligands in the initial and selected structures, representing substantial diversity in the bound ligand. If only one crystal structure was available for the isoform, additional structural models were created withAlphaFold3 and RoseTTAFold2 [16,17]. The protein descriptors were calculated using the iFeature Python package, with 65 protein sequence descriptors and 14 protein structure descriptors [18]

**Table 1.** Selected 3D Structures of Cytochrome P450 Isoforms for Model Training and Testing

|  |  |  |  |
| --- | --- | --- | --- |
| Isoform | PDB code (Highest Resolution) | PDB code (Most different by pocket RMSD) | PDB code (Most different by ligand Tanimoto Score) |
| Training Set |  |  |  |
| CYP2C9 | 5A5I | 1OG5 | 6VLT |
| CYP2C19 | 4GQS | * | # |
| CYP2D6 | 3TBG | 4XRZ | 3QM4 |
| CYP3A4 | 4D6Z | 2J0D | 8EXB |
| External Validation Set |  |  |  |
| CYP17A1 | 6WR1 | 3SWZ | 8FDA |
| CYP1A2 | 2HI4 | * | # |
| The table outlines the selected 3D structures from the RCSB Protein Data Bank (PDB) for each cytochrome isoform used in the study. For each isoform, three conformations were identified: the structure with the highest resolution, the structure with the greatest backbone variation (measured by pocket RMSD compared to the highest resolution structure), and the structure representing the most distinct active site composition based on the Tanimoto distance between ligands. * - structures generated using AlphaFold-3, # - structures generated using RoseTTAFold2. |  |  |  |

### Dataset collection and processing: ligand selection

The binding ligands were extracted from the ChEMBL database, using a cutoff value of 1 µM for Ki and IC50 (see Table 2). Structures containing salts and duplicate entries between the training and test sets were identified and excluded. Due to the inherent imbalance in the binder/non-binder ratio within the ChEMBL database, resulting from the limited availability of reported non-binders, a corresponding set of DUD-E decoys was generated [19]. The dataset was organized in tabular format with binary class assignments for each data point. Ligand molecules were stored in the canonical SMILES format. 3D conformations were generated, and polar hydrogens were added at pH 7.4 using Schrodinger LigPrep.

**Table 2.** Summary of Dataset Structure for Cytochrome P450-Ligand Binding Models

|  |  |  |  |
| --- | --- | --- | --- |
| Isoform | No. of ligands before clustering | No. of ligands after clustering | No. of decoys after clustering |
| Training set |  |  |  |
| CYP2C9 | 529 | 204 | 10200 |
| CYP2C19 | 253 | 149 | 7450 |
| CYP2D6 | 723 | 299 | 14950 |
| CYP3A4 | 1300 | 414 | 20700 |
| Test set |  |  |  |
| CYP1A2 | 534 | 217 | 10850 |
| CYP17A1 | 738 | 174 | 8700 |
| The dataset includes the number of binding ligands before clustering, the number of DUD-E decoys generated for each protein and the number of ligands retained after clustering. The training set comprises four proteins (CYP2C9, CYP2C19, CYP2D6, and CYP3A4), while the external testing set includes CYP17A1 and CYP1A2. Ligands were clustered using the DBSCAN method with a Tanimoto distance threshold of 0.4 to ensure structural diversity in the validation sets. |  |  |  |

### Computation of Ligand Molecular Descriptors

A set of 156 physical and chemical molecular descriptors was computed using KNIME software v. 4.18.0 using the RDKit, Indigo, and CDK packages [20,21,22,23] (see Supplementary Table 4). Topological coordinates descriptors were not included in the analysis since their use could result in intentional distinguishing between active compounds and DUD-E decoys, as these decoy molecules are specifically generated with distinct topological features while maintaining physical similarity to active compounds. SMILES molecules were converted into Morgan fingerprints for these calculations [24]. The resulting molecular descriptors were stored in tabular format for each CYP450-ligand pair, with each descriptor calculated across all three selected isoforms. In cases where only one protein structure was available for a given isoform, the descriptor values were duplicated to preserve the consistency of the tabular structure.

### Docking calculations and rescoring

Molecular docking calculations were performed using the Glide software suite in standard precision mode (Schrödinger, LLC, v. 2024-3) [25]. The Protein Preparation Wizard module was utilized to preprocess all protein structures. The protein structures of the CYP450 isoforms were prepared by removing water molecules, adding hydrogen atoms, and assigning partial charges using the OPLS3 force field. Ligands, including both experimentally validated binders and non-binders, were prepared using LigPrep to generate possible ionization states and tautomers. The receptor grid was initially prepared using the Receptor Grid Generation Panel in the Schrodinger suite, with a cubic inner box (15 Å per side) and an outer box (20 Å per side), centered at the crystal ligand from the initial pdb structure. Following docking, the results were analyzed to assess ligand binding poses and scored to evaluate binding affinity. GlideScore, an empirical scoring function which includes force field terms and rewarding/penalizing terms, was used to evaluate the docking results. Individual components of scoring functions were also analysed individually (Supplementary Table 4).

To evaluate whether changing the scoring function would affect the predictive model’s accuracy, we used scoring functions (SFs) of different classes. Scoring functions can be broadly categorized into physics-based, knowledge-based, and empirical SFs, each utilizing distinct principles to estimate binding [26]. In this study, ITScoreAff was chosen as a representative knowledge-based SF. ITScoreAff is trained on both structural and affinity data of 6216 protein-ligand complexes [27]. Cyscore, an empirical SF, estimates the protein-ligand binding free energy by decomposing it into four key components: hydrophobic free energy, van der Waals interaction energy, hydrogen bond interaction energy, and the ligand’s conformational entropy [12]. By including these two SFs, we aimed to assess their influence on docking performance and to diversify the output of molecular docking. The obtained results from rescoring were added to hybrid models via individual descriptors as described in Supplementary Table 4.

### Protein-ligand interaction descriptors calculation

SIFt (Structural Interaction Fingerprint) is a method that converts three-dimensional protein-ligand interaction information into a one-dimensional binary string (fingerprint), enabling efficient analysis, visualization, and comparison of binding patterns in protein-ligand complexes [28]. In this work, SIFts were calculated using the Schrödinger Suite from the docking poses obtained through Glide docking [25]. The extracted SIFts were subsequently transformed into interaction descriptors, capturing key features of the protein-ligand interactions, such as hydrogen bonds, hydrophobic contacts, and π-π stacking interactions (Supplementary Table 4). These interaction descriptors were then used as input features for the predictive models. By incorporating SIFt-based descriptors, we aimed to improve the model’s ability to distinguish between binders and non-binders, particularly by providing detailed information on interaction patterns that ligand descriptors might miss.

### Machine Learning Framework

The proposed model utilized a stacked ensemble approach, implemented through the AutoGluon-Tabular classifier within a machine learning framework [29]. The AutoGluon-Tabular module incorporates a set of baseline ML algorithms, including K -Nearest Neighbors, tree-based methods (Random Tree and Random Forest), boosting methods (LightGBM, XGBoost, and CatBoost), a weighted ensemble method, and neural network-based approaches (NeuralNet and NeuralNetTorch). AutoGluon automates the hyperparameter tuning process, streamlining model optimization and achieving enhanced performance via multi-stacking.

The machine learning models were trained with a train-to-test ratio of 80:20 and 50-fold validation on a combined dataset of 4 CYP450 isoforms (CYP2C9, CYP2C19, CYP2D6, CYP3A4). In this work models with the highest logAUC score were extracted to compare the performance of the individual top scored models with the Weighted Ensemble model. A weighted ensemble is a machine learning method in which individual models within an ensemble are given weights according to their performance metrics. The final prediction is generated by aggregating the predictions of these models through a weighted average or voting mechanism, giving greater influence on more accurate models, allowing more accurate models to have a greater impact on the final prediction. The ensemble then combines predictions using a weighted voting or averaging technique. AutoGluon employs a weighted ensemble method that combines individual model predictions into a single output through a weighted sum. The assigned weights remain consistent across all time series within the dataset.

AutoGluon employs a k-fold cross-validation strategy, where the dataset is split into K subsets, and N models are trained iteratively on K-1 folds while reserving one for validation, obtaining N x K models in total. With the last layer of ensemble model, AutoGluon adds a “meta-model” to the equation, resulting in M x N x K + 1 models, where the resulting ensemble “meta-model”. Model iterations are trained on different data partitions, generating out-of-fold (OOF) predictions for evaluation. The final model performance is assessed using OOF predictions against ground truth labels. During inference, AutoGluon aggregates predictions from all trained models, averaging them, to improve accuracy and generalization. This method ensures robust model selection while minimizing overfitting.

In this work, we divided the machine learning framework into two distinct models: Ligand Model and Hybrid Model, as illustrated in Figure 1. Each model was trained on a different set of descriptors. Specifically, the Ligand Model utilized only ligand descriptors, while the Hybrid Model combined ligand, protein and protein-ligand interaction descriptors (including molecular docking, rescoring and SIFt). This division allowed us to evaluate the individual and cumulative contributions of each descriptor type to the predictive performance of CYP450 binding. We conducted 50-fold validation, ensuring data was randomly shuffled for each iteration to enhance the reliability of the results. The outcomes were also compared to molecular docking calculations performed using Glide.

Furthermore, we assessed the models by considering the use of multiple protein structures. For each CYP450 isoform, the model was evaluated using the primary protein structure with the highest resolution available, along with two additional protein structures/models. These additional structures were selected to capture the structural diversity of the binding site, ensuring that the model could generalize its predictions across multiple protein structures. This approach allowed us to understand the impact of protein structure variability on model performance and improve the robustness of our predictions for CYP450-ligand binding.

### Predictive Model Analysis

The performance of these methods was evaluated using a range of accuracy metrics. The logAUC metric, which represents ligand enrichment and serves as the primary accuracy evaluation metric in this work, is frequently used for benchmarking molecular docking calculations, which is helpful for comparing the results of this work to the current state-of-art methods [10].

In addition to logAUC, we also incorporated AUPR (Area Under the Precision-Recall Curve) and AUC score to gain a more comprehensive evaluation of model performance, particularly in imbalanced datasets. The AUPR metric focuses on the precision and recall of the model, providing a more informative assessment in scenarios where the dataset contains a significant imbalance between positive and negative classes. It emphasizes the model’s ability to correctly identify positive samples without being biased by the overwhelming presence of negative samples. The AUC (Area Under the ROC Curve) indicates the likelihood that the model will assign a higher score to a randomly selected positive instance than to a randomly selected negative one. The Area Under the ROC Curve (AUC) is a widely used evaluation metric in machine learning, which is particularly commonly reported in the literature on CYP450-ligand prediction. However, it is more appropriate for balanced datasets, as it does not account for class imbalance and may present an overly optimistic view of model performance in imbalanced scenarios.

### External Testing and Benchmarking with State-of-Art Models

The external testing was conducted on an independent dataset, including the CYP1A2 and CYP17A1 isoforms, to assess the generalizability of the models to novel cytochrome P450 targets. For external testing, we chose the Weighted Ensemble Multiple Structure Hybrid Model that demonstrated the highest predictive performance and the Weighted Ensemble Ligand Model as a reference to analyze how the inclusion of protein and protein-ligand interaction descriptors can affect the generalizability of the model. Additionally, models were compared against molecular docking results to evaluate the relative performance of external testing.

The benchmarking analysis was conducted for CYP3A4 and CYP1A2 isoforms and was performed using the Hybrid Ensemble Multi-Structure Model that was described previously. This model was benchmarked against two state-of-art CYP450-ligand binding prediction tools: SwissADME and ADMETlab 3.0 [13,14]. Both tools are based on ligand descriptors, without explicitly considering protein structural variability. The comparison focused on three key evaluation metrics: AUPR (Area Under the Precision-Recall Curve), and AUC (Area Under the Curve) score. The benchmarking was conducted using binders and non-binders from the ChEMBL database, described after the releases of SwissADME and ADMETlab 3.0 models in respective articles. To ensure diversity, the ligands were clustered using the Butina method with a Tanimoto similarity threshold of 0.6, and the cluster centroids were chosen for benchmarking.

## ASSOCIATED CONTENT

The Supporting Information is available free of charge.

Molecular docking results represented by logAUC values. (Table S1); 80/20 validation results for CYP2D6, CYP3A4, CYP2C19, CYP2C9 represented by logAUC value (Table S2); External validation results for CYP1A2 and CYP17A1 isoforms represented by logAUC values (Table S3); List of descriptors used in Ligand and Hybrid models sorted by class categories (Table S4); Relative descriptor importance in the Weighted Ensemble Multiple Structure Hybrid Model for CYP450 binding prediction represented by normalized importance values (Table S5); Benchmarking results for CYP1A2 and CYP3A4 isoforms represented by AUC and AUPR values (Table S6).

## ACKNOWLEDGEMENT

This project is supported by the Singapore International Graduate Award (SINGA), A*STAR, Singapore (to A.T.), and Bioinformatics Institute, A*STAR, Singapore (to H.F.). The Table of Contents illustration is created with BioRender.com (Tikhonova, A. (n.d.) https://BioRender.com/yn8pg8g)

## DATA AVAILABILITY STATEMENT

The list of molecules used in this model is provided in ChEMBL_molecules.xlsx file in https://github.com/aetikh/cyp450_hybrid/. This repository also contains a tutorial on setting up and running the model described in the article in CYP450_Hybrid.ipynb. The list of required molecular descriptors is provided in Supplementary_Information.docx

## ABBREVIATIONS

AUC: Area Under the Curve
AUPR: Area Under the Precision-Recall Curve
CDK: Chemistry Development Kit
ChEMBL: Chemical Biology Database
CYP: Cytochrome P450
DUD-E: Directory of Useful Decoys - Enhanced
HTS: High-Throughput Screening
IC50: Half-maximal Inhibitory Concentration
ITScoreAff: Information Theory Scoring Function for Affinity
logAUC: Logarithmic Area Under the Curve
MD: Molecular Dynamics
PDB: Protein Data Bank
PLEC: Protein-Ligand Extended Connectivity
RMSD: Root Mean Square Deviation
SIFt: Structural Interaction Fingerprint
SMILES: Simplified Molecular Input Line Entry System

## For Table of Contents Only

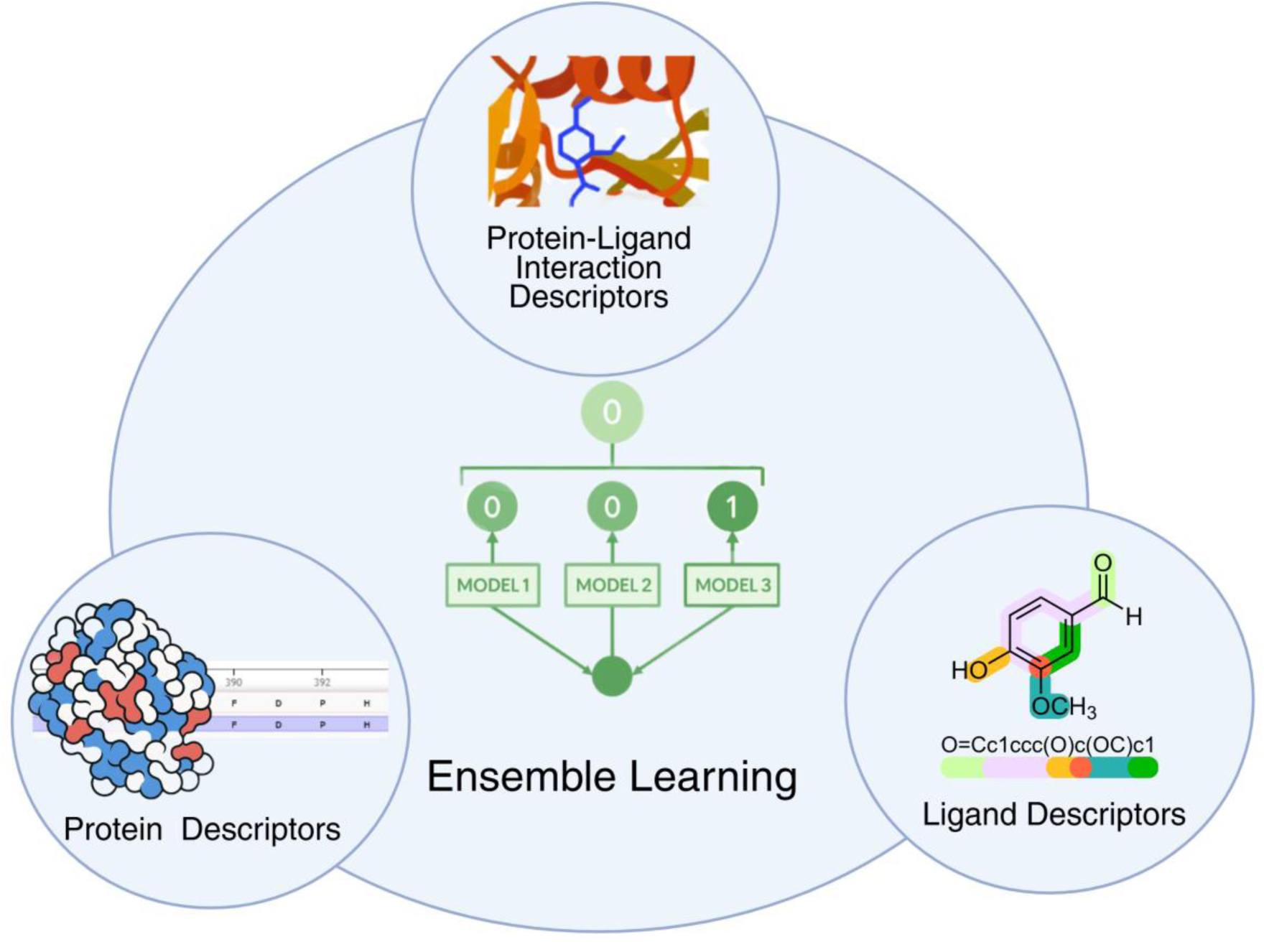
Created in BioRender. Tikhonova, A. (n.d.) https://BioRender.com/yn8pg8g

## Notes

### Competing Interest Statement

The authors have declared no competing interest.

