## Supplementary Information for "Enhancing CYP450-Ligand Binding Predictions: A Comparative Analysis of Ligand-Based and Hybrid Machine Learning Models"

| Supp. Table 1. Molecular docking results represented by logAUC values. | | | |
|---|---|---|---|
| Protein | PDB | logAUC | Consensus logAUC* |
| CYP2C9 | 5A5I | 0.24 | 0.3 |
| | 1OG5 | 0.26 | |
| | 6VLT | 0.24 | |
| CYP2C19 | 4GQS | 0.24 | 0.35 |
| | * | 0.36 | |
| | # | 0.23 | |
| CYP2D6 | 3TBG | 0.24 | 0.26 |
| | 4XRZ | 0.24 | |
| | 3QM4 | 0.23 | |
| CYP3A4 | 4D6Z | 0.24 | 0.25 |
| | 2J0D | 0.25 | |
| | 8EXB | 0.24 | |
| CYP1A2 | 2HI4 | 0.20 | 0.28 |
| | * | 0.26 | |
| | # | 0.22 | |
| CYP17A1 | 6WR1 | 0.24 | 0.25 |
| | 3SWZ | 0.24 | |
| | 8FDA | 0.23 | |
| *Consensus logAUC is calculated by selecting the most negative docking scores among 3 presented protein structures. * - structures generated using AlphaFold-3, # - structures generated using RoseTTAFold2. | | | |

Supp. Table 2. 80/20 validation results for CYP2D6, CYP3A4, CYP2C19, CYP2C9 represented by logAUC values.

| Method | CYP2C9 | CYP2C19 | CYP2D6 | CYP3A4 |
|---|---|---|---|---|
| Top Scored Ligand Model | 0.40 ± 0.022 | 0.38 ± 0.016 | 0.45 ± 0.043 | 0.38 ± 0.023 |
| Weighted Ensemble Ligand Model | 0.43 ± 0.027 | 0.41 ± 0.012 | 0.49 ± 0.062 | 0.39 ± 0.022 |
| Top Scored Single Structure Hybrid Model | 0.90±0.020 | 0.81 ± 0.039 | 0.9±0.059 | 0.97±0.074 |
| Weighted Ensemble Single Structure Hybrid Model | 0.92±0.015 | 0.82±0.038 | 0.92±0.029 | 0.98±0.048 |
| Top Scored Multiple Structure Hybrid Model | 0.94±0.078 | 0.84±0.071 | 0.93±0.121 | 0.98±0.003 |
| Weighted Ensemble Multiple Structure Hybrid Model | 0.95±0.074 | 0.86±0.131 | 0.96±0.183 | 0.99±0.004 |

Error values indicate standard deviation across 50-fold cross-validation

| Supp. Table 3. External validation results for CYP1A2 and CYP17A1 isoforms represented by logAUC values. | | |
|---|---|---|
| Method | CYP1A2 | CYP17A1 |
| Top Scored Ligand Model | 0.28 | 0.47 |
| Weighted Ensemble Ligand Model | 0.3 | 0.49 |
| Top Scored Single Structure Hybrid Model | 0.31 | 0.95 |
| Weighted Ensemble Single Structure Hybrid Model | 0.36 | 0.97 |

| Supp. Table 4. List of descriptors used in Ligand and Hybrid models sorted by class categories. | |
|---|---|
| Physico-chemical descriptors | SP3 Character, Molar Mass, Formal Charge, Formal Charge (pos), Formal Charge (neg), XLogP, Vertex adjacency information magnitude, Topological Polar Surface Area, Lipinski's Rule of Five, Largest Chain, VABC Volume Descriptors, Bond Count, Bond Polarizabilities, Element Count, Aromatic Bonds Count, Aromatic Atoms Count, Atomic Polarizabilities, Mannhold LogP, Number of heteroatoms in aliphatic rings, Number of chiral centers, Number of connected components, Number of heteroatoms, Number of cis/trans bonds, Number of R-sites, Number of hydrogens, Total number of atoms, Number of heteroatoms in aromatic rings, Number of aromatic rings, Number of bonds, Number of visible atoms, Number of aliphatic atoms, Number of aliphatic bonds, NumAliphaticCarbocycles,NumSaturatedCarbocycles,NumAromaticCarbocycles, NumSaturatedHeterocycles, NumAliphaticRings, NumSaturatedRings, NumRings, NumStereocenters, NumAmideBonds, NumLipinskiHBD, NumLipinskiHBA, LabuteASA,SMR, Molecular Quantum Number Descriptors (MQN1-MQN42), Subdivided Surface Area Descriptors (SlogPVSA0-SlogPVSA9, SMR_VSA0-SMR_VSA7), Partial Equalization of Orbital Electronegativities (PEOE_VSA+6-PEOE_VSA-6). |
| Protein descriptors | Amino acids content type 1 (AAC_type1), Amino acids content type 2 (AAC_type2), Grouped amino acids content type 1 (GAAC_type1), Grouped amino acids content type 2 (GAAC_type2), Secondary structure elements (3) type 1 (SS3_type1), Secondary structure elements (3) type 2 (SS3_type2), Secondary structure elements (8) type 1 (SS8_type1), Secondary structure elements (8) type 2 (SS8_type2), Half sphere exposure α (HSE_CA), Half sphere exposure β (HSE_CB), Residue depth (Residue depth), Atom content type 1 (AC_type1), Atom content type 2 (AC_type2), Network-based index |
| SIFT descriptors | SIFT Charged Residue, SIFT Aromatic Residue, SIFT Hydrogen Bond Donor, SIFT Hydrogen Bond Acceptor, SIFT Hydrophobic Residues, SIFT Polar Residues, SIFT Sidechain Interaction, SIFT Backbone Interaction,SIFT Any Contact |
| Docking and Rescoring descriptors | Glide Rotatable Bonds, Glide Einternal, Glide Energy, Glide Score, Glide Emodel, Glide Ecoul, Glide Evdw, Glide hbond, Glide lipo, ITScoreAff, Cyscore,Cyscore_Ent, Cyscore_HBond, Cyscore_Vdw,Cyscore_Hydrophobic, Cyscore,Cyscore_Ent, Cyscore_HBond, Cyscore_Vdw, Cyscore_Hydrophobic |

| Supp. Table 5. Relative descriptor importance in the Weighted Ensemble Multiple Structure Hybrid Model for CYP450 binding prediction represented by normalized importance values. | |
|---|---|
| Feature | Value |
| Glide Docking Score | 0.31 |
| ITScore | 0.3 |
| SIFT Charged Residue | 0.12 |
| Cyscore_Vdw | 0.11 |
| Cyscore_HBond | 0.033 |
| Cyscore_Hydrophobic | 0.031 |
| SIFT Aromatic Residue | 0.03 |
| SIFT Sidechain Interaction | 0.022 |
| MQN17 | 0.02 |
| MQN31 | 0.02 |

| Suppl. Table 6. Benchmarking results for CYP1A2 and CYP3A4 isoforms represented by AUC and AUPR values. | | | | |
|---|---|---|---|---|
| Model | CYP3A4 | | CYP1A2 | |
| | AUC | AUPR | AUC | AUPR |
| ADMET 3.0 | 0.47 | 0.51 | 0.44 | 0.47 |
| SwissADME | 0.51 | 0.53 | 0.56 | 0.54 |
| Hybrid Model | 0.76 | 0.78 | 0.61 | 0.57 |
